# Benchmarking supervised signature-scoring methods for single-cell RNA sequencing data in cancer

**DOI:** 10.1101/2021.06.29.450404

**Authors:** Nighat Noureen, Zhenqing Ye, Yidong Chen, Xiaojing Wang, Siyuan Zheng

## Abstract

Quantifying the activity of gene expression signatures is common in analyses of single-cell RNA sequencing data. Methods originally developed for bulk samples are often used for this purpose without accounting for contextual differences between bulk and single-cell data. More broadly, these methods have not been benchmarked. Here we benchmark four such supervised methods, including single sample gene set enrichment analysis (ssGSEA), AUCell, Single Cell Signature Explorer (SCSE), and a new method we developed, Jointly Assessing Signature Mean and Inferring Enrichment (JASMINE). Using cancer as an example, we show cancer cells consistently express more genes than normal cells. This imbalance leads to bias in performance by bulk-sample-based ssGSEA in gold standard tests and down sampling experiments. In contrast, single-cell-based methods are less susceptible. Our results suggest caution should be exercised when using bulk-sample-based methods in single-cell data analyses, and cellular contexts should be taken into consideration when designing benchmarking strategies.

## Background

Gene signatures are an important tool for making sense of omics datasets. Historically defined as sets of genes that display coordinated expression changes upon perturbation, this definition has been expanded to include biochemical pathways, genes residing in the same genomic locations, co-regulated targets, or any sets that share certain connections^1^. Gene lists resulting from omics datasets, typically by comparing two phenotypes, are often tested for signature enrichment. The enrichment results provide interpretability and insights into the contrasting phenotypes. Although this approach has been highly successful, one caveat is the rapid increase of analytic complexity when more than two phenotypes are involved.

An alternative class of approaches alleviates this complexity by quantifying signature activities in individual samples without using information from other samples. Outputs from these supervised methods can thus be conveniently correlated with complex sample groupings. More importantly, they provide a tool to interrogate dynamic biological properties such as sternness and telomerase activity in cancers^2,3^ that are otherwise challenging to quantify for large cohorts.

Single-cell RNA sequencing (scRNA-seq) is a powerful technology to study cellular heterogeneity in normal tissues and diseases^4,5^. After cell clusters are identified from scRNA-seq, they are annotated for cell identity and states based on single markers or marker signatures. Supervised methods are well suited for scoring marker gene signatures. With more and more cell types being discovered and characterized, this usage can be further extended to match a cell population to a reference cell atlas as a generic approach complementary to more sophisticated methods^6,7^.

For these reasons, supervised signature-scoring methods have been increasingly used in scRNA-seq data analysis. A widely used method is single sample gene set enrichment analysis (ssGSEA)^8–18^, which was developed for bulk samples^19^. However, bulk-sample RNAseq is distinct from scRNA-seq in that the latter has much higher variability, particularly high dropout rates^20^. The dropout rate, or conversely the number of genes detected, was recently associated with cell differentiation status^18^. Thus, in contexts where cells differ in differentiation status, using this method may lead to inaccurate results. More broadly, few attempts have been made to benchmark these methods.

Cancer cells exhibit stem-cell like properties^3^, which are generally lacking in normal cells. In this work, we benchmark four signature-scoring methods in cancer, including ssGSEA (implemented in the R GSVA package^19^) and three single-cell-based methods, AUCell^21^, SingleCell Signature Explorer (SCSE)^22^, and Jointly Assessing Signature Mean and INferring Enrichment (JASMINE), a new method we developed. We initially included other bulk-sample-based methods such as GSVA, but it was much slower in computational speed thus was dropped. We show that cancer cells consistently express more genes (we use the number of expressed genes and gene counts interchangeably hereafter) than normal cells, and this imbalance significantly biases results from ssGSEA but largely spares single-cell-based methods. Though this benchmarking study was done in cancer datasets, in practice, we used these datasets only to recapitulate the variability in gene counts. Our results caution against the use of bulk-sample-based methods in scRNA-seq analyses.

## Results

### Cancer cells demonstrate higher gene counts than normal cells

Several recent studies showed that gene counts of single cells are associated with cell differentiation status^18^ and cell identity^23,24^. Specifically, cells higher in the differentiation hierarchy express more genes. Cancer cells display stem cell features compared to most normal cells from the same tissue of origin^25^. To examine if they demonstrate the same bias, we collected 10 scRNA-seq datasets across different cancer types, technological platforms and sequencing coverage (**Supplementary Table 1**). We also obtained cell type annotations from the original studies. We found that the average number of detected genes was significantly higher in tumor cells than in normal cells across all datasets (**Fig. 1a**, p-value < 2.2e-16; **Supplementary Table 1**). This imbalance persisted when we separated normal cells into different cell populations (**Supplementary Fig. 1**). However, cancer-associated cells, including cancer-associated fibroblasts (CAF), tumor-associated macrophages (TAM) and tumor-related endothelial cells (TEC) had higher gene counts than other normal cell types, sometimes even comparable to malignant cells. (**Supplementary Fig. 1**).

**Fig 1.**
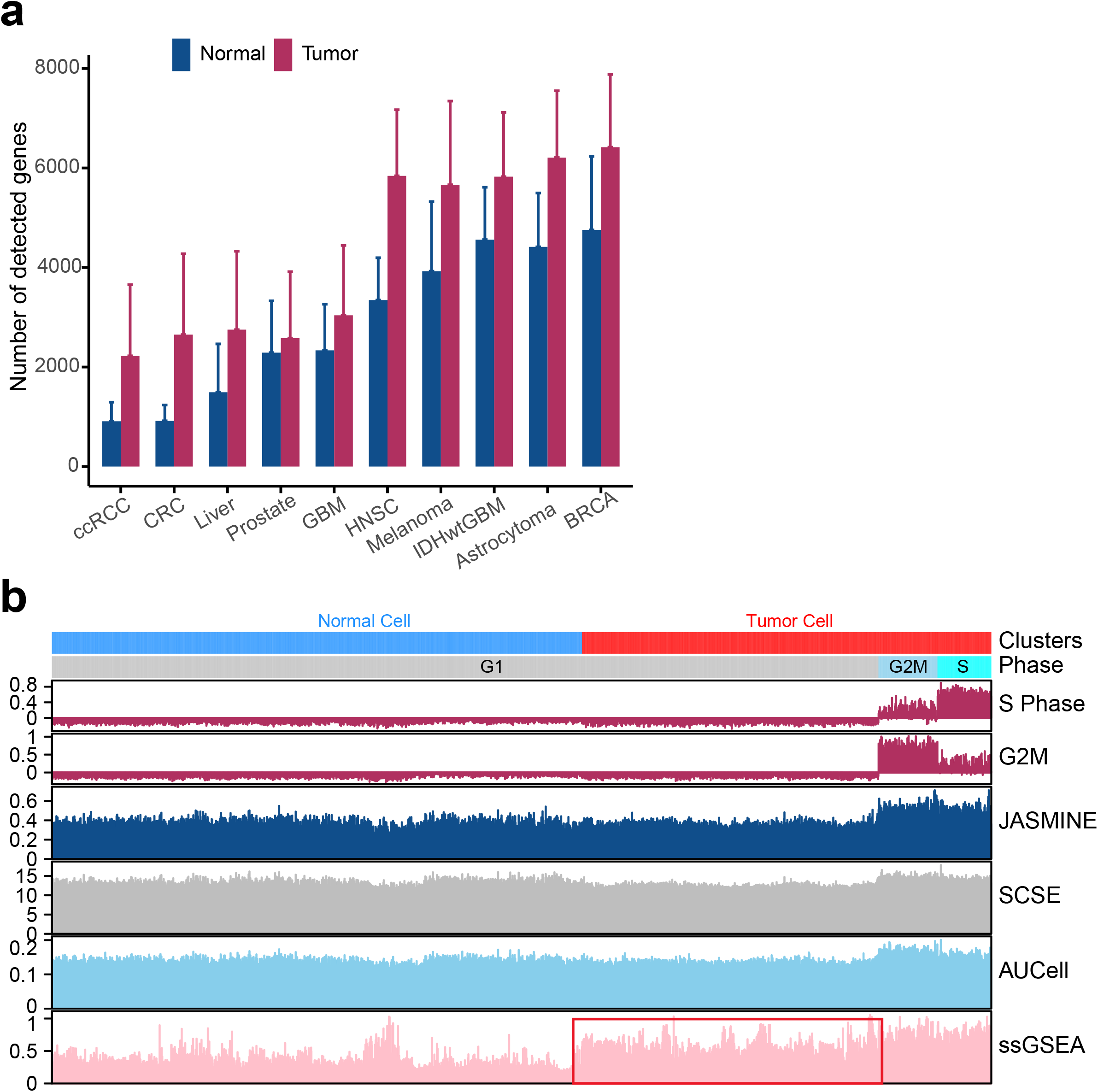
Gene counts imbalances affect signature scoring. **(a)** The number of detected genes in tumor and normal cell populations in 10 cancer single cell RNAseq data sets. More details of these datasets can be found in Supplementary Table 1. In all cases, the difference is statistically significant (student t test, p<2.2e-16). **(b)** An example of cell cycle scores from Seurat compared with Gene Ontology cell cycle signature (GO:0007049) scored using JASMINE, AUCell, SCSE and ssGSEA in tumor and normal cell populations. Cell cycle phases are divided into cycling (S, G2M) and non-cycling (G1). Head and Neck cancer dataset is used for this example. Red box highlights non-cycling cancer cells that received higher scores than non-cycling normal cells.

To illustrate the potential impact of gene counts on signature scoring, we scored the cell cycle gene signature from Gene Ontology^1^ (GO:0007049) using the head and neck cancer (HNSC) data set^26^. This signature contains genes regulating G2M and S phases, and thus, should confer a higher score to cycling cells. As a benchmark, we used Seurat^27^ to identify cycling cells at G2M and S phases. As expected, all four methods reported a higher score in cycling cells **(Fig. 1b)**. However, ssGSEA also reported higher scores in non-cycling cancer cells than normal cells although neither are cycling. In contrast, the single-cell-based methods did not show this difference. These results imply that ssGSEA may inadequately account for data variability in scRNA-seq data.

### Bias in ssGSEA signature scores in cancer datasets

We next sought to systematically benchmark these methods. Because of their quantitative nature, we reasoned that the reliability of such methods lies in their ability to identify differences, or no differences, between two group of samples at the gene-signature level. One challenge is that scores from these methods differ in range and variance. To overcome this challenge, we used Cohen’s d^28^, a measure of effect size commonly used in meta-analysis. Cohen’s d normalizes mean differences with standard deviation, thus generating unitless contrast that can be compared across methods. For simplicity, we defined Cohen’s d greater than 1 as up regulated, and less than −1 as down regulated. These criteria were used throughout this manuscript unless otherwise noted.

We assembled 7600 gene sets that each has at least 20 genes (Methods). Three out of the 10 scRNA-seq datasets were dropped because they lacked the similar gene count bias at the signature level **(Supplementary Fig. 2)**. The remaining datasets were then used to score all gene sets using the four methods.

We first compared tumor and normal cells **(Fig. 2a)**. On average, single-cell methods reported similar numbers of up and down regulated gene sets (13% vs 11%), except in a few datasets AUCell and SCSE reported more down gene sets. In contrast, ssGSEA reported 36% of the input gene sets as up regulated and 6% as down regulated. Specifically, in six out of seven datasets, ssGSEA estimated more than twice as many up regulated gene sets than down gene sets. This pattern remained when we categorized normal cells into cell populations **(Fig. 2b–2c and Supplementary Fig. 3)**. Notably, in cell types with high gene counts such as TEC, TAM and CAF, we did not observe significantly more up regulated gene sets in ssGSEA **(Supplementary Fig. 3c)**. These results collectively suggest ssGSEA is sensitive to variability of gene counts common to scRNA-seq data.

**Fig 2.**
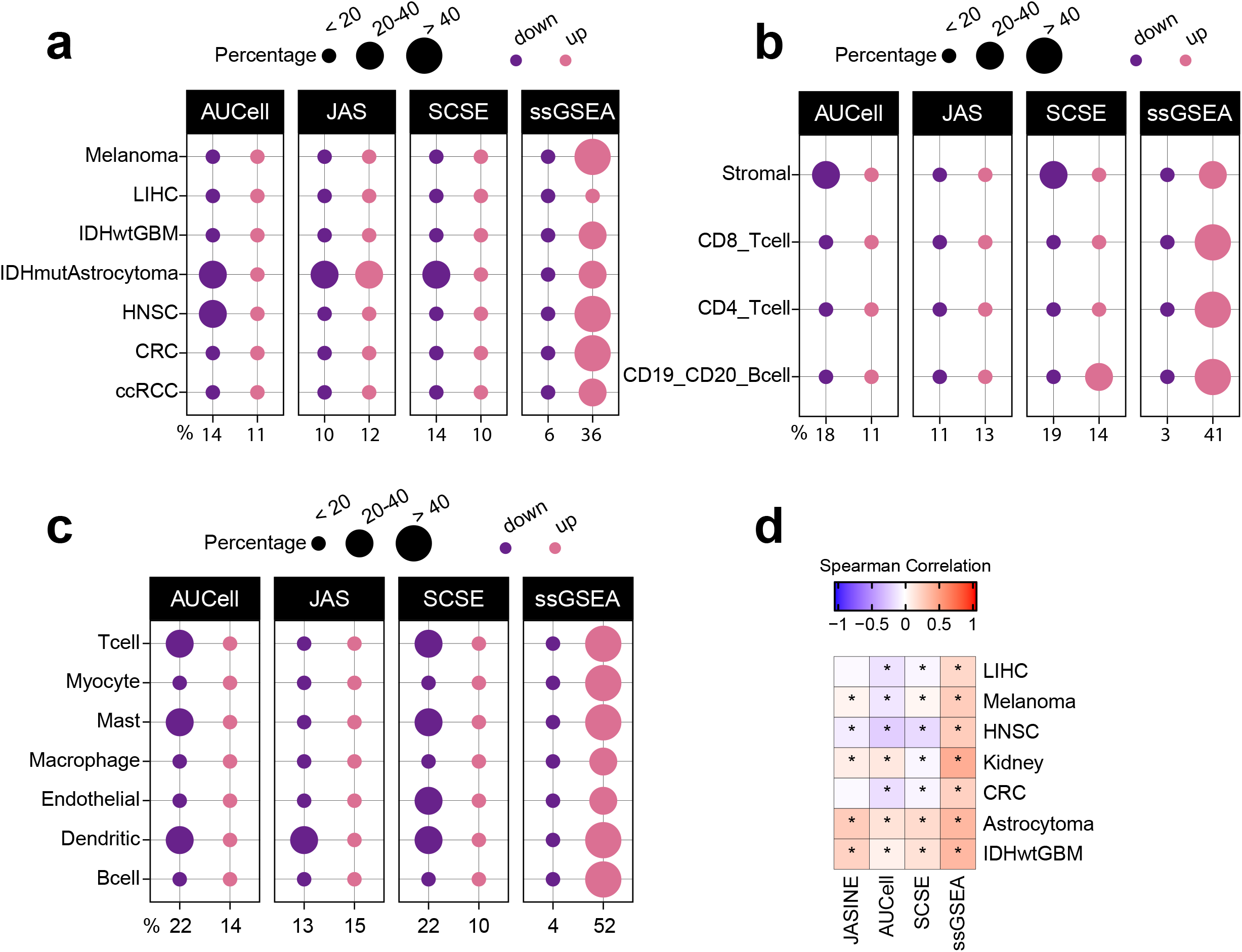
Overall scoring pattern of the four tools in cancer and normal cell comparison. (a) Percentage of up and down regulated gene signatures in cancer cells relative to normal cells based on Cohen’s d. Dot size corresponds to the percentage of all signatures tested (n=7600). (b) Comparison between cancer cells and different normal cell populations using the colorectal dataset. (c) Comparison between cancer cells and different normal cell populations using the head and neck cancer dataset. (d) Spearman correlation coefficients of Cohen’s d with signature sizes across the datasets and methods. Asterisk (*) in each cell indicates p-value < 0.01. Color of the heatmap represents correlation coefficient.

We also found that Cohen’s d from ssGSEA showed consistent positive correlations with gene set sizes **(Fig. 2d)**. Single-cell methods did not demonstrate this pattern.

### Benchmarking detection sensitivity

We next used simulation to generate gold standard gene sets to benchmark detection sensitivity for these methods. We first identified differentially expressed genes for each dataset using MAST^29^, a method that explicitly accounts for dropout rates. We then randomly drew *N* genes from up regulated genes to generate an up gene set of size *N*, and similarly for down gene sets. Because in practice up and down regulated gene sets contain genes that have no expression change, or even changes at opposite directions, we added noises to the simulated gene sets. For instance, when setting noise level to 20%, an up gene set of size *N* would have 20% of its genes drawn from the remainders other than the up regulated genes. Following this procedure, we generated gene sets at the sizes of 50, 100, 150, 200, and 300 genes. For each size, we set the noise levels to 0, 20%, 40%, 60% and 80%. For each noise-size combination, we randomly generated 200 gene sets and used the average to represent the setting.

**Fig. 3** summarizes the results for both up and down simulated gene sets. A more detailed illustration is provided in **Supplementary Fig. 4.** We found gene set size had negligible impacts on detection rates. Noise levels, on the other hand, had a much bigger impact. For up gene sets, all methods were able to detect 50% of the gene sets on average even at 80% noise level **(Fig. 3a).** However, for down gene sets, ssGSEA performed worse than other methods (**Fig. 3b**). Without noise, it only detected about 80% of the gene sets. At 80% noise level, it detected around 30% of gene sets, compared to 70%-80% by single-cell methods.

**Fig 3.**
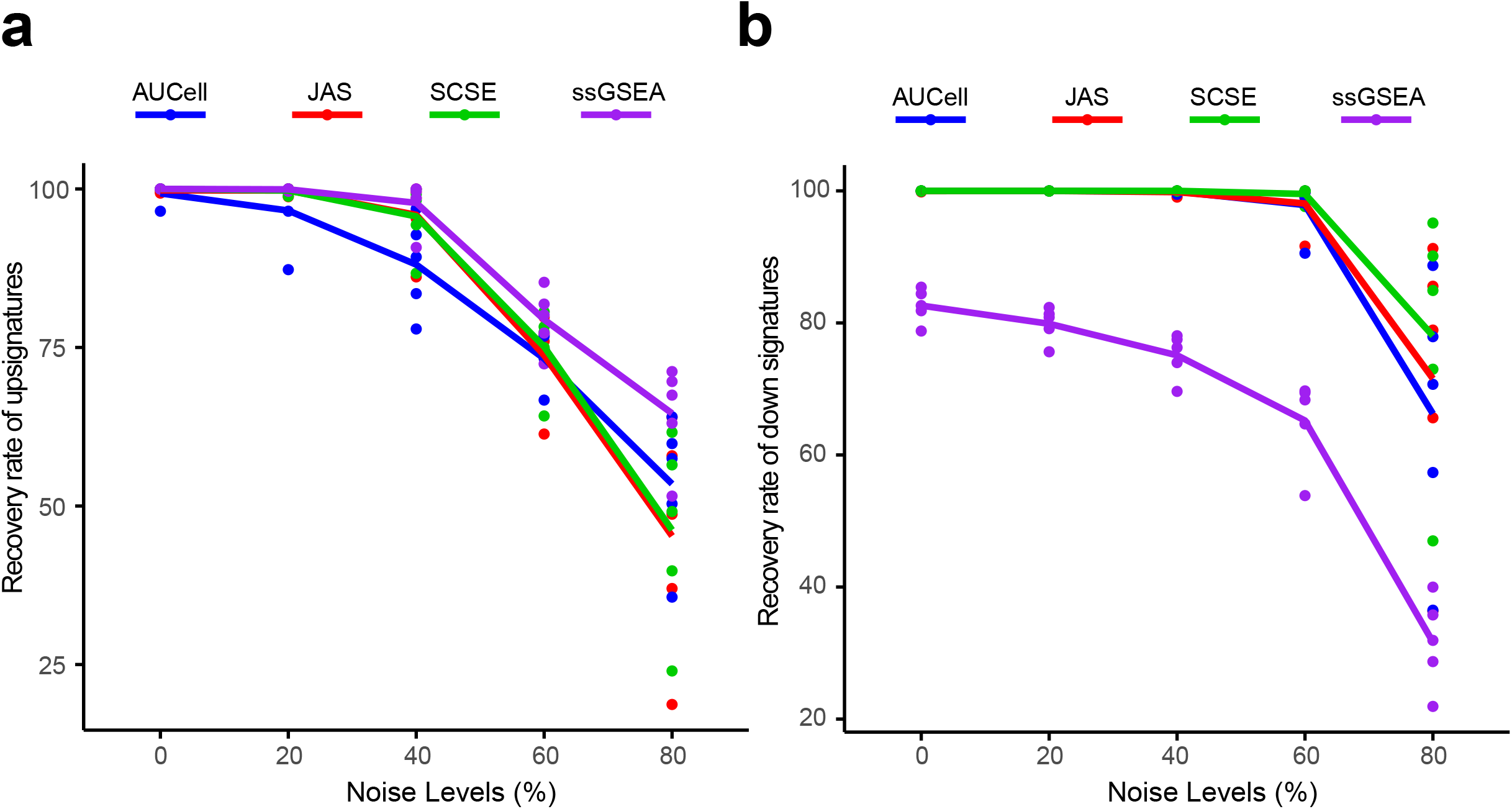
Recovery rates for simulated up and down signatures. **(a)** Recovery rate for up gene signatures across 5 noise levels by the four methods. Each dot represents one dataset. At each noise level, average of all datasets is used to represent the performance of each method. **(b)** Similarly for down signatures.

### Benchmarking detection specificity

To benchmark detection specificity, we generated null datasets by randomly choosing 100 cancer cells from each dataset. We down sampled each cell to 50% coverage, resulting in lower gene counts. These cells were then normalized to the same total coverage. Because the down-sampled cells were the same cells with the original, no up or down gene sets were expected between the two groups.

We found AUCell and JASMINE outperformed SCSE and ssGSEA in detection specificity **(Fig. 4a)**. On average, AUCell detected 11% and 2% of the input gene sets as up and down; JASMINE detected 6% and 2%. In two datasets, SCSE overestimated down gene sets (average 19%). ssGSEA grossly overestimated both up and down gene sets in all datasets (average 16% for up and 37% for down).

**Fig 4.**
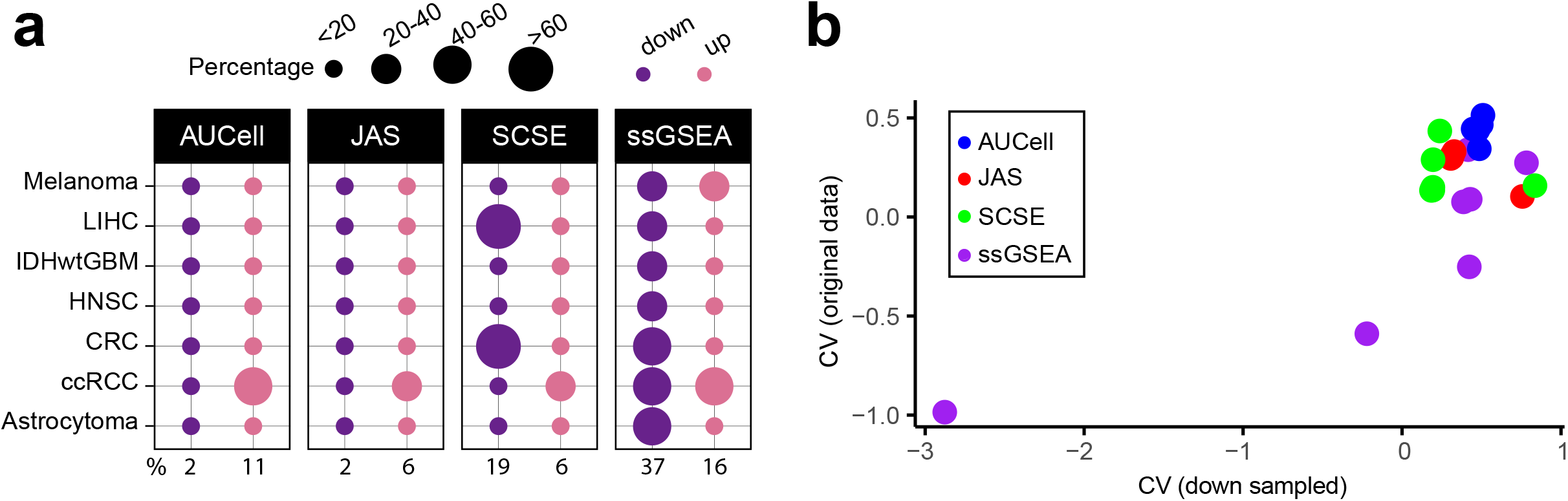
False discovery estimates based on down sampling. **(a)** Percentages of false up and down signatures. The size of the dots corresponds to the percentages of all the signatures tested. Because the contrasting groups are generated by down sampling, no signatures are expected to be identified. The numbers below the heatmap are the average percentage. **(b)** Average coefficient of variance between the original datasets and the 50% down-sampled datasets. Each dot represents one dataset.

Next, we estimated how changes in gene count affected score stability. For each gene signature, we calculated coefficient of variance (CV) **(Fig. 4b)**. We observed that single-cell methods were very robust to our down sampling, with little changes in CV. However, in three datasets, ssGSEA had a higher CV in down-sampled cells than in original cells.

### Comparing single-cell methods to consensus

Our benchmarking analysis using simulation data show that single-cell methods generally performed well. We next compared the three methods against consensus—gene sets that were identified as up or down by at least two methods. Though consensus is not ground truth, it is more reliable than individual methods, as was shown in many benchmark studies^30,31^.

We observed a better consistency between JASMINE and the consensus reflected by its higher accuracy in most datasets **(Fig. 5a and Supplementary Fig. 5).** AUCell overall had lower accuracies, particularly for down gene sets, presumably because it is designed to score marker signatures **(Fig. 5a).** When breaking down to sensitivity and specificity, AUCell showed higher false positive rates **(Supplementary Fig. 6a).** The three tools showed comparable sensitivity. Across the seven datasets, SCSE and JASMINE showed a better correlation **(Fig. 5b** and **Supplementary Fig. 6b)**, thus explaining their relatively better alignment with the consensus.

**Fig 5.**
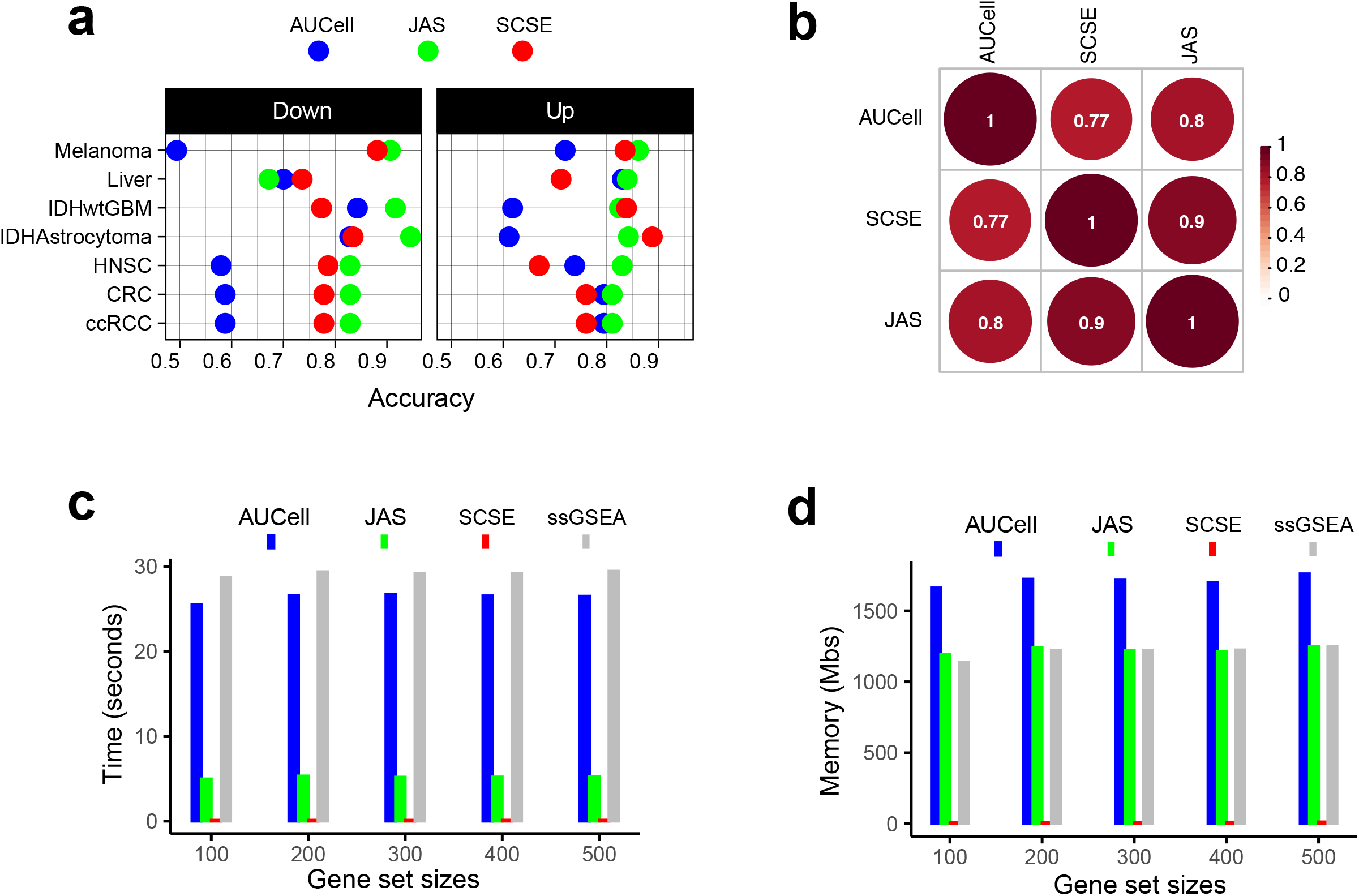
Benchmarking single-cell methods against consensus and evaluation of computing cost. **(a)** Accuracy of the three methods, separated into up and down signatures. Accuracy is calculated as the agreement with consensus calls by at least two methods. **(b)** Consistency between the three methods. Numbers are Spearman correlation coefficients. **(c)** Average time consumption for completing 50 gene signatures using a 2.2 GHz, 32 GB memory CPU. (d) Memory cost for completing 50 gene signatures using a 2.2 GHz, 32 GB memory CPU.

Finally, we tested computing efficiency of the three methods. We randomly generated gene sets of various sizes and analyzed each gene set in a dataset consisting of 2,000 cells. **Fig. 5c** and **5d** show the time and memory use averaged over 50 iterations for one gene set under the same hardware configuration (2.20 GHz CPU, 32 GB memory). SCSE took least time and memory. JASMINE was the second fastest algorithm, but its memory use was same as that of ssGSEA. AUCell was slightly faster than ssGSEA but required more memory. Their performances were robust across different signature sizes.

## Discussion

In this study, we benchmarked four non-parametric signature-scoring methods in seven cancer scRNA-seq datasets. Based on overall pattern, comparison with gold standard gene sets, and down sampling, we show that single-cell-based methods are more robust at identifying *bona fide* up and down gene signatures. In contrast, bulk-sample-based ssGSEA is more susceptible to gene count variability and computationally more expensive. Among single-cell methods, JASMINE and SCSE showed better concordance, and JASMINE outperforms SCSE in down sampling tests.

One limitation of this study is that we only tested cancer datasets. Other contexts may also bear the same gene count imbalance, such as in stem cells versus mature cells. However, cell identities are largely irrelevant in our testing other than their association with gene counts. In addition, generating gold standard gene sets in our study involved drawing genes from a pool of up or down genes identified by an established algorithm^29^. Since no algorithm performs perfectly, this pool may include false positives that can confound gene set simulation. Alternatively, we could artificially generate up and down genes by purposely altering their expression values in a selected group of cells and then draw gene sets from these genes. This approach rigorously establishes gold-standard differentially expressed genes, as was shown in related benchmark experiments^32^, and thus, will minimize the effects of false positive genes. However, it ignores the differences in cellular contexts that may be associated with cells’ global expression patterns by changing the expression of only a few genes while leaving others unchanged.

In conclusion, our results caution against using bulk-sample-based signature-scoring methods to score single cells, which often vary in cellular contexts. In contrast to single cells, cellular contexts of bulk samples are often blurred due to their mixture nature. As more and more single cell states and types are discovered, the full scale of the impacts of cellular contexts on gene expression will be better understood and will in return used to refine analytic methods. Our study also suggests that it is important to consider cellular contexts when benchmarking methods for scRNA-seq data analysis, particularly when bulk-sample-based methods are involved.

## Methods

### Single cell data sets and Signature scoring

We used single cell data sets from 10 published studies^26,33–41^ for the evaluation of number of expressed genes in tumor versus normal cells to identify significant heterogenous patterns among the two phenotypes. We used C2 (n=6,226), C3 (n=3,556) and Hallmarks (n=50) modules from Msigdb^1^ v.7.2, to calculate the ratio of signature genes across all data sets and further signature scoring. Based on the significant differential pattern of signature genes across 2 phenotypes **(Supp. Fig. 2),** we shortlisted 7^26,33–36,38,39^ out of 10 data sets for signature score calculations using SCSE^22^, AUCell^21^, ssGSEA^19^ and JASMINE.

### Jointly Assessing Signature Mean and Inferring Enrichment (JASMINE)

For each signature, JASMINE calculates the approximate mean using gene ranks among expressed genes and the enrichment of the signature in expressed genes. The two are then scaled and averaged to result in the final JASMINE score.

Assume *R_gc_* represents the rank of an expressed signature gene *g* among genes with expression value > 0 in a cell *c*, then a mean rank vector *V_mean_* would be as follows:

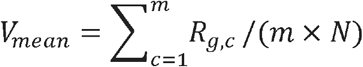

Where *m* represents the total number of expressed genes in a signature and *N* represents the total number of expressed genes per cell. Because the mean is based on ranks, it is robust to scale normalization methods. It assesses the expression level of the signature only using genes detected in one cell, thus minimizing the effect of dropouts.

To assess signature enrichment in the expressed genes, we calculate Odds Ratio (OR) using 4 variables: a = signature genes expressed, b = signature genes not expressed, c = non-signature genes expressed and d = non-signature genes not expressed. Then signature enrichment (OR) is calculated as follows:

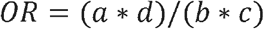

In practice, *c* is unlikely zero. For smaller signatures, b can be occasionally 0. In that case, we replace it with 1. OR assesses the distribution of signature genes against the dropout background. Finally, *V_mean_* and *OR* are linearly scaled to [0 - 1] and are averaged to generate JASMINE scores.

### Effect size for gene sets scores

We first filtered the 9,835 gene sets to 7,503 by requiring a minimum size more than 20. We scored 7600 gene sets (C2, C3 and hallmarks) among tumor and normal phenotypes using Cohen’s d^28^. We utilized an R package “effectsize”^42^ to calculate this metric. We used Cohen’s d >= 1 as a threshold for positive or up cases and <= −1 as negative or down cases with respect to tumor versus normal cells.

### Gene signature simulation

To generate gold standard gene signatures, we first identified differentially expressed genes using MAST^29^ (hurdle model with default settings). To generate an up-gene signature that contains 50 genes at the noise level x, we randomly selected 50*x genes from the up regulated genes, and selected 50*(1-x) genes from the other genes including down regulated genes and genes that do not show differential expression. This process was repeated for 200 times to generate 200 such signatures. Similar procedures were used to generate signatures at sizes 100, 150, 200 and 300 at noise levels 0, 20%, 40%, 60% and 80% for both up and down directions. In total, 5000 gene sets were generated per data set for each direction. All four methods were utilized to score these gene sets.

### Down sampling

We subset 100 tumor cells per data set for down sampling. The R package “scuttle”^43^ was used to down sample each cell to 50% total coverage. The down sampled data sets were then scaled back to ensure equal total coverage before comparison.

## Data Availability

Single cell data sets used in this study including their downloading sources were listed in **Supplementary Table 1.** Gene sets were downloaded from MSigDB v.7.2 (http://www.gsea-msigdb.org/gsea/msigdb/index.jsp) for C2, C3 and Hallmark modules. JASMINE source code is available on Github (https://github.com/NNoureen/JASMINE).

---

**Supplementary Table 1. Datasets used in the study.** Details of the 10 datasets evaluated, including source, size, platform, annotation etc.

**Supplementary Fig 1. Bias in gene counts.** Number of genes expressed in tumor and normal cell types across 7 single cell data sets including Colorectal cancer (CRC), Liver cancer (LIHC), Head and Neck Cancer (HNSC), Melanoma and Clear Cell Renal Carcinoma (ccRCC). Maroon colors represent tumor cell populations in each data set while blue represents normal cell populations across all data sets. Note that in the LIHC dataset, CAF, TAM, TEC are cancer-associated fibroblasts, tumor-associated macrophages, and tumor-related endothelial cells.

**Supplementary Fig 2. Bias in gene counts at signature level.** Each column is one signature (total n=7600), and each row represents one dataset. Color represents log2(Tumor/Normal) in terms of expressed genes in the signature. A few datasets, including GBM, BRCA, Prostate, do not show significant bias in signature gene counts between tumor and normal, even though the overall gene counts differ. Therefore, these three datasets were dropped from our analysis.

**Supplementary Fig 3. Patterns of up and down regulated signatures comparing tumor and normal cell** populations across four additional datasets, including (a) astrocytoma, (b) IDHwt GBM (c) Liver, and (d) Melanoma. The size of each dot represents the percentage of up or down signatures over all signatures tested (n=7600).

**Supplementary Fig 4. Benchmarking sensitivity using simulated gene signatures.** We simulated four gene set sizes (50,100,150,200 and 300), each with five levels of noise (0, 20%, 40%, 60% and 80%). For each size/noise combination, we randomly generated 1000 signatures. The results shown in this figure are average of the 1000 random signatures. **(a)** Detection sensitivity for up gene signatures. Deeper color indicates lower recovery rates (thus more misses). **(b)** detection sensitivity for down signatures.

**Supplementary Fig 5. Comparison of calling results from the four methods across the seven datasets.** In heatmap, each column represents one signature. Blue, down signature; Red, up signature.

**Supplementary Fig 6. Consistency with consensus and pairwise comparison. (a)** Sensitivity and false positive benchmarked against the consensus calls (signatures called by at least two methods). (b) Spearman correlation of Cohen’s d broken down to each dataset.

