## Supplementary Figure 1 for "Benchmarking supervised signature-scoring methods for single-cell RNA sequencing data in cancer"

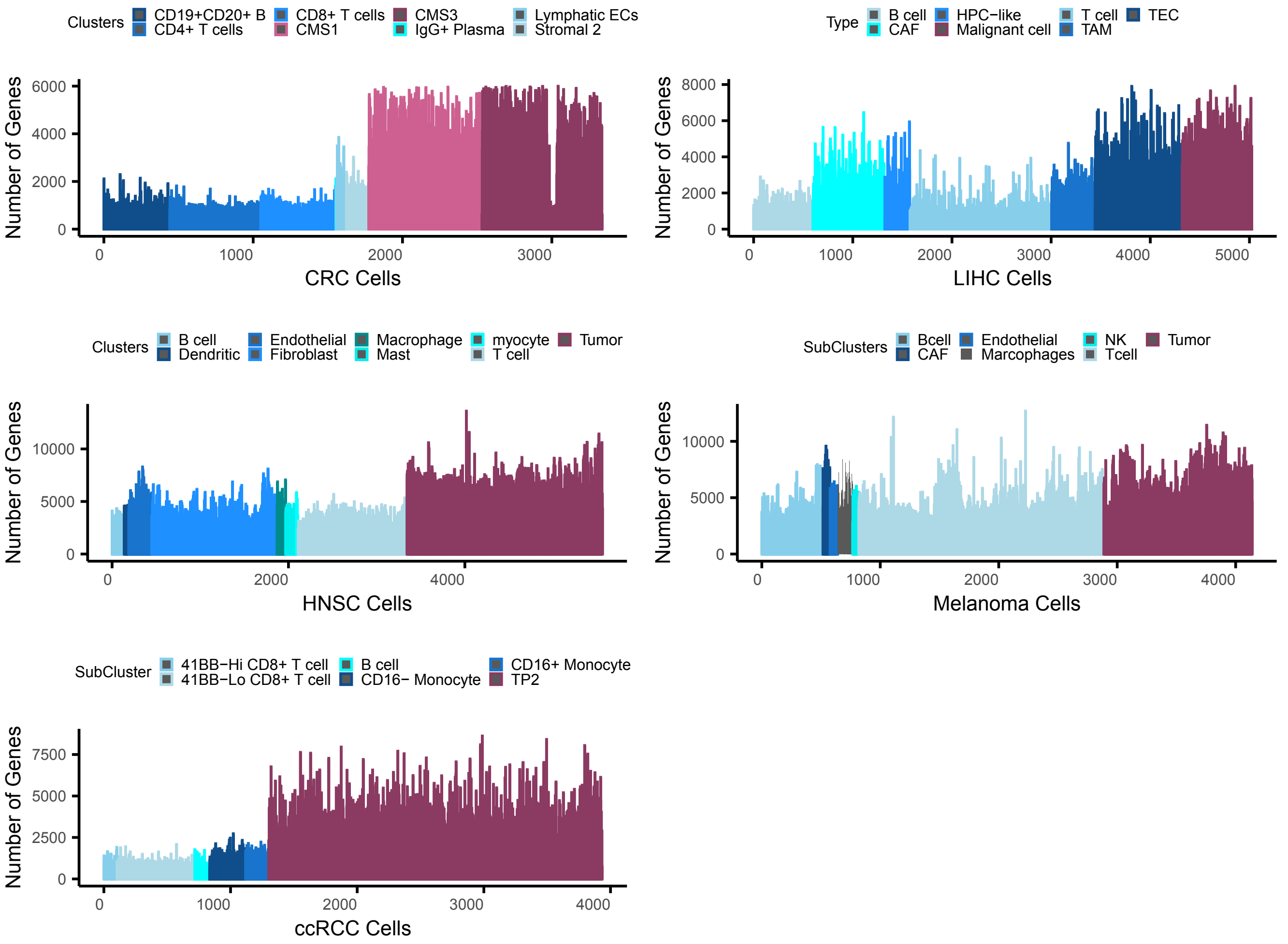


**Supplementary Fig 1. Bias in gene counts.** Number of genes expressed in tumor and normal cell types across 7 single cell data sets including Colorectal cancer (CRC), Liver cancer (LIHC), Head and Neck Cancer (HNSC), Melanoma and Clear Cell Renal Carcinoma (ccRCC). Maroon colors represent tumor cell populations in each data set while blue represents normal cell populations across all data sets. Note that in the LIHC dataset, CAF, TAM, TEC are cancer-associated fibroblasts, tumor-associated macrophages, and tumor-related endothelial cells.
