## Supplementary Figure 2 for "Benchmarking supervised signature-scoring methods for single-cell RNA sequencing data in cancer"

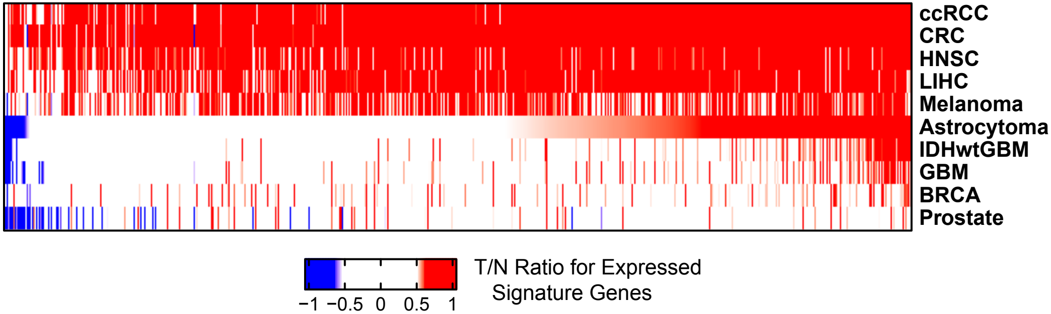


**Supplementary Fig 2. Bias in gene counts at signature level.** Each column is one signature (total n=7600), and each row represents one dataset. Color represents log2(Tumor/Normal) in terms of expressed genes in the signature. A few datasets, including GBM, BRCA, Prostate, do not show significant bias in signature gene counts between tumor and normal, even though the overall gene counts differ. Therefore, these three datasets were dropped from our analysis.
