## Supplementary Figure 3 for "Benchmarking supervised signature-scoring methods for single-cell RNA sequencing data in cancer"

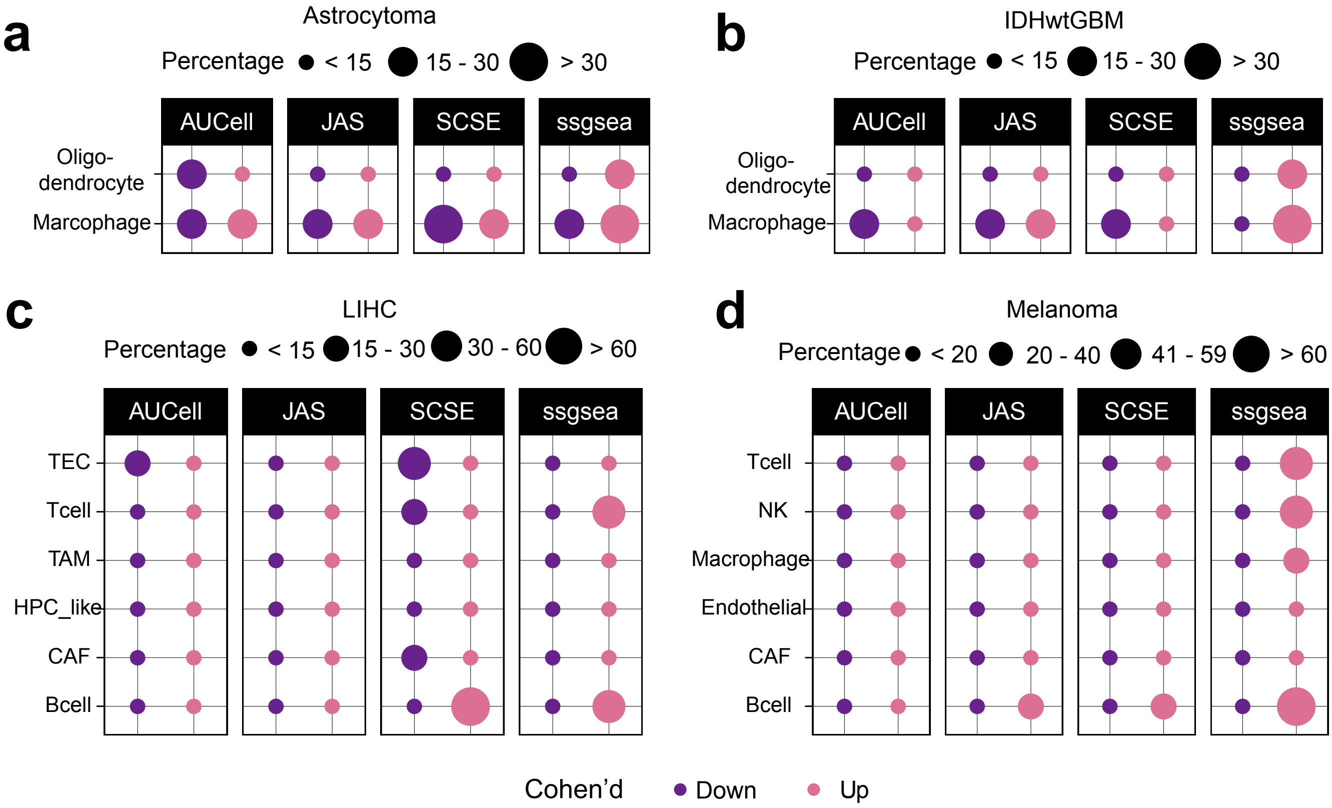


**Supplementary Fig 3. Patterns of up and down regulated signatures comparing tumor and normal cell populations** across four additional datasets, including (a) astrocytoma, (b) IDHwt GBM (c) Liver, and (d) Melanoma. The size of each dot represents the percentage of up or down signatures over all signatures tested (n=7600).
