## Supplementary Figure 4 for "Benchmarking supervised signature-scoring methods for single-cell RNA sequencing data in cancer"

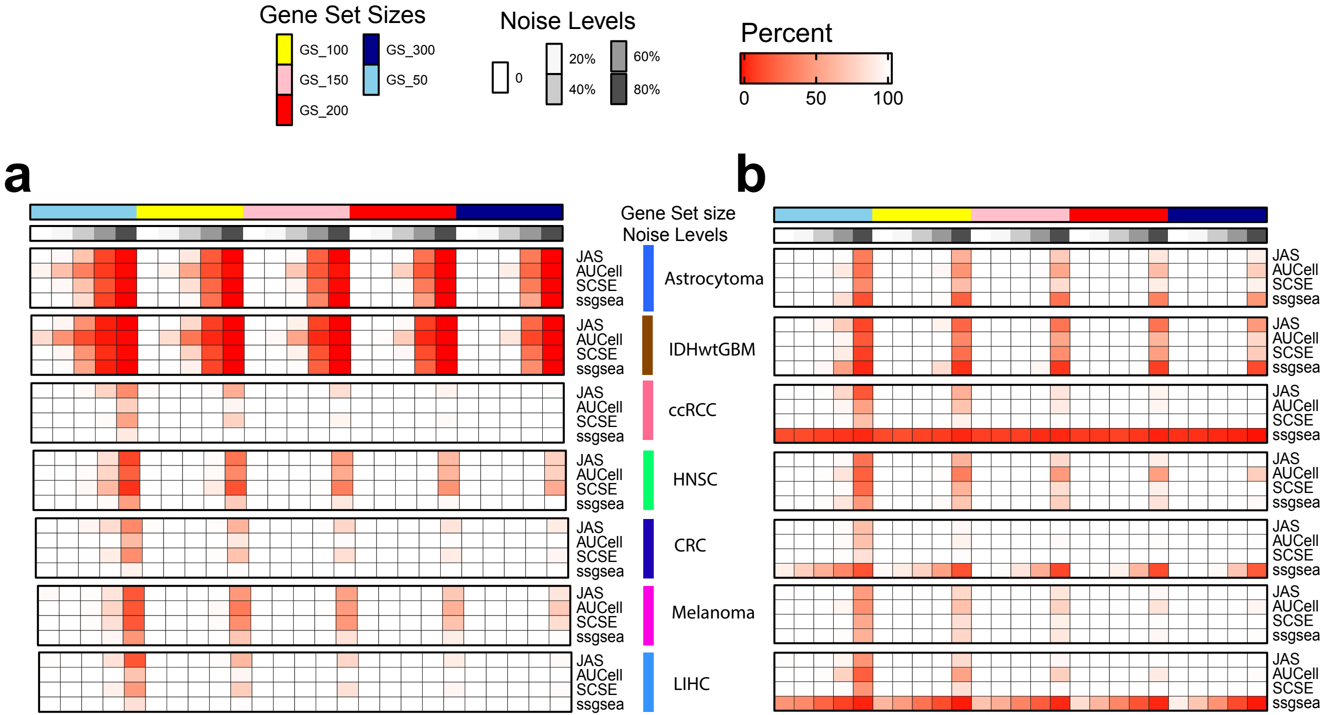


**Supplementary Fig 4. Benchmarking sensitivity using simulated gene signatures.** We simulated four gene set sizes (50,100,150,200 and 300), each with five levels of noise (0, 20%, 40%, 60% and 80%). For each size/noise combination, we randomly generated 1000 signatures. The results shown in this figure are average of the 1000 random signatures.
**(a)** Detection sensitivity for up gene signatures. Deeper color indicates lower recovery rates (thus more misses). **(b)** detection sensitivity for down signatures.
