## Supplementary Figure 5 for "Benchmarking supervised signature-scoring methods for single-cell RNA sequencing data in cancer"

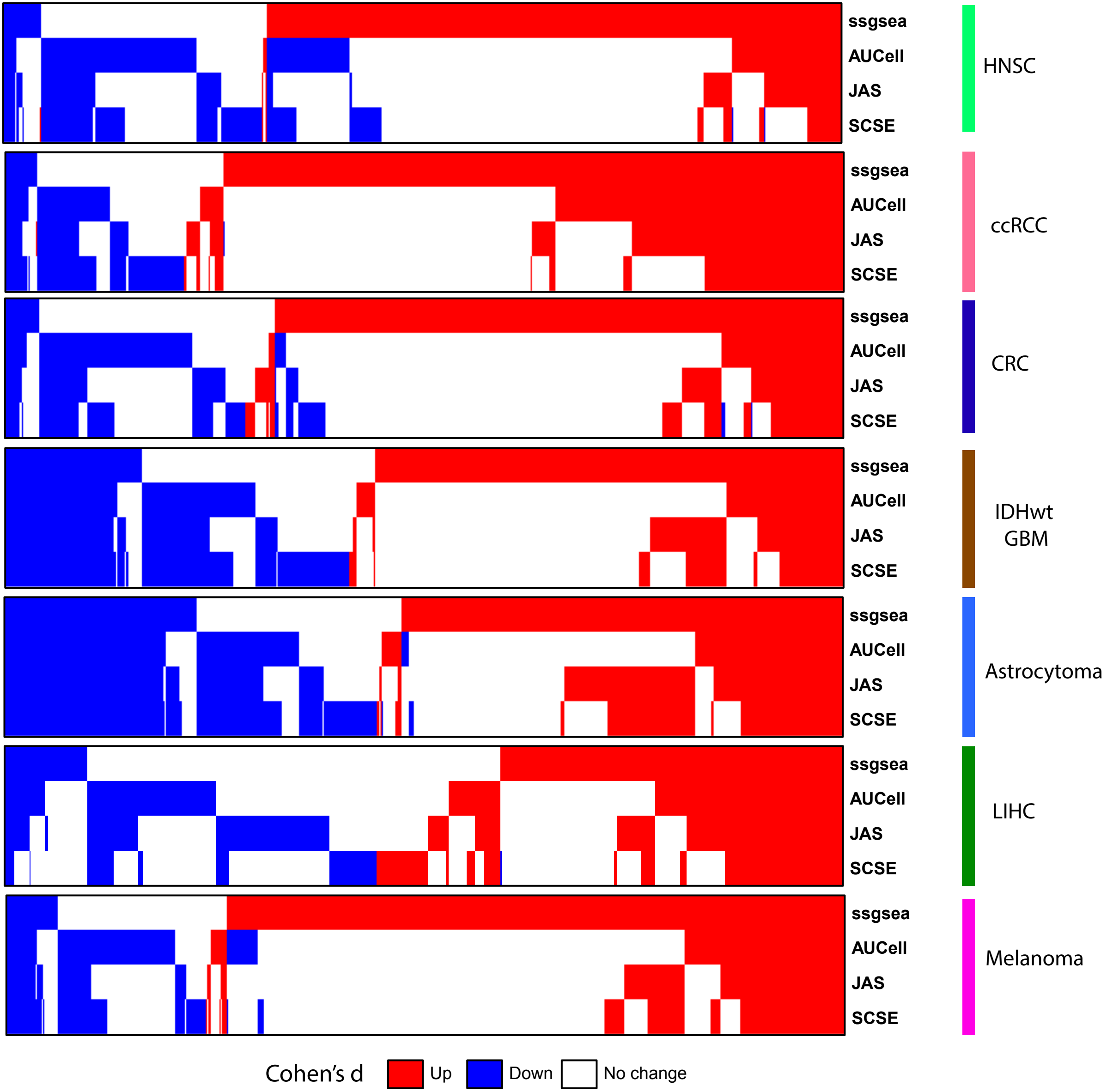


**Supplementary Fig.5. Comparison of calling results from the four methods across the seven datasets.** In heatmap, each column represents one signature. Blue, down signature; Red, up signature.
