## Supplementary Figure 6 for "Benchmarking supervised signature-scoring methods for single-cell RNA sequencing data in cancer"

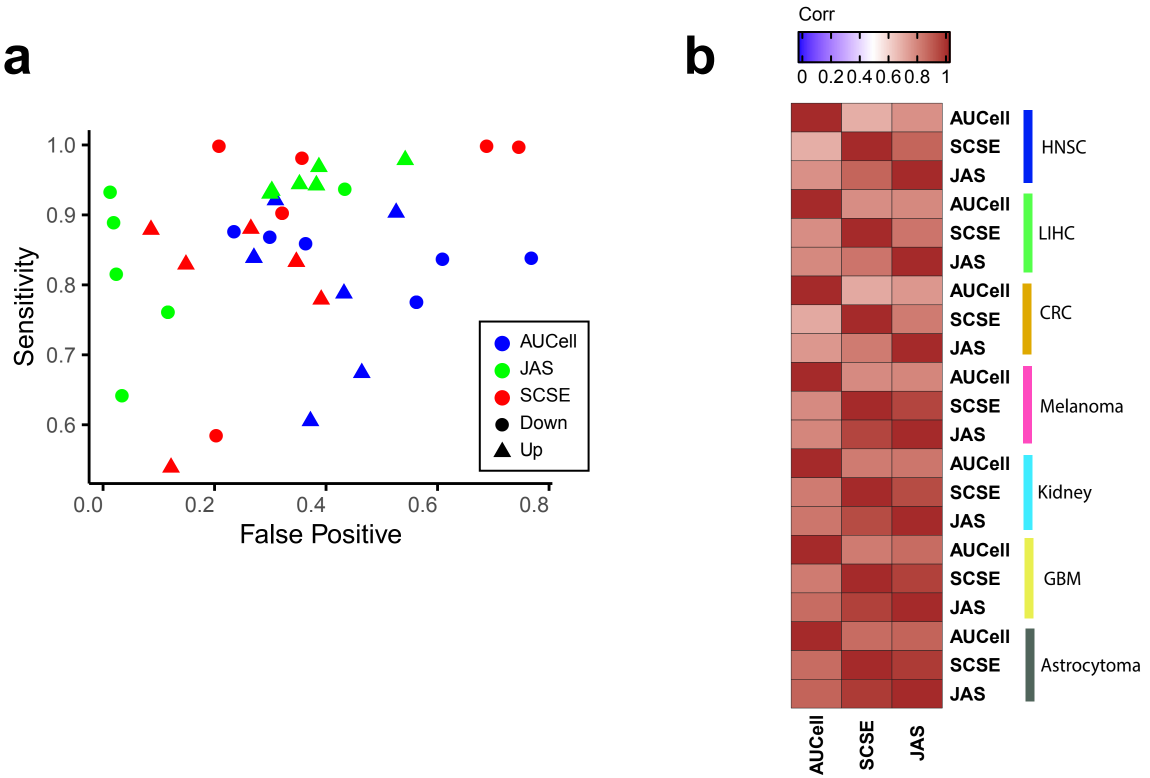


**Supplementary Fig 6. Consistency with consensus and pairwise comparison. (a)** Sensitivity and false positive benchmarked against the consensus calls (signatures called by at least two methods). **(b)** Spearman correlation of Cohen's d broken down to each dataset.
